# Planning horizon affects prophylactic decision-making and epidemic dynamics

**DOI:** 10.1101/069013

**Authors:** Luis G. Nardin, Craig R. Miller, Benjamin J. Ridenhour, Stephen M. Krone, Paul Joyce, Bert O. Baumgaertner

## Abstract

Human behavior can change the spread of infectious disease. There is limited understanding of how the time in the future over which individuals make a behavioral decision, their planning horizon, affects epidemic dynamics. We developed an agent-based model (along with an ODE analog) to explore the decision-making of self-interested individuals on adopting prophylactic behavior. The decision-making process incorporates prophylaxis efficacy and disease prevalence with individuals' payoffs and planning horizon. Our results show that for short and long planning horizons individuals do not consider engaging in prophylactic behavior. In contrast, individuals adopt prophylactic behavior when considering intermediate planning horizons. Such adoption, however, is not always monotonically associated with the prevalence of the disease, depending on the perceived protection efficacy and the disease parameters. Adoption of prophylactic behavior reduces the peak size while prolonging the epidemic and potentially generates secondary waves of infection. These effects can be made stronger by increasing the behavioral decision frequency or distorting an individual’s perceived risk of infection.

## Introduction

Human behavior plays a significant role in the dynamics of infectious disease [7, 11]. However, the inclusion of behavior in epidemiological modeling introduces numerous complications and involves fields of research outside the biological sciences, including psychology, philosophy, sociology, and economics. Areas of research that incorporates human behavior into epidemiological models are loosely referred to as *social epidemiology, behavioral epidemiology*, or *economic epidemiology*[14, 17]. We use the term ‘behavioral epidemiology’ to broadly refer to all epidemiological approaches that incorporate human behavior. While the incorporation of behavior faces many challenges [9], one of the goals of behavioral epidemiology is to understand how social and behavioral factors affect the dynamics of infectious disease epidemics. This goal is usually accomplished by coupling models of social behavior and decision making with biological models of contagion [16, 11].

Many social and behavioral aspects can be incorporated into a model of infectious disease. One example is the effect of either awareness or fear spreading through a population [10, 5]. In these types of models, the spread of beliefs or information is treated as a contagion much like an infectious disease, though the network for the spread of information may differ from the biological network [2]. Other models focus on how individuals adapt their behavior by weighting the risk of infection with the cost of social distancing [6, 19] or other disincentives [1]. Still others model public health interventions (e.g. isolation, vaccination, surveillance, etc.) and individual responses to them[3]. Many of these models sit at the population level, incorporating the effects of social factors and abstracting away details about the individuals themselves.

The SPIR model (**S**usceptible, **P**rophylactic, **I**nfectious, **R**ecovered) is an epidemiological agent-based model that couples individual behavioral decisions with an extension of the SIR model [13]. In this model agents that are vulnerable to infection may be in one of two states which are determined by their behavior. Agents in the susceptible state engage in the status quo behavior while agents in the prophylactic state employ preventative behaviors that reduce their chance of infection. We use a rational choice model to represent individual behavioral decisions, where individuals select the largest utility between engaging in prophylactic behavior (e.g. hand-washing or wearing a face mask) or non-prophylactic behavior (akin to the status quo). We also allow for the fact that individuals may not perceive the risk of getting infected accurately, but rather receive some distorted information, for example, through the media.

We are interested in understanding how an individual’s planning horizon—the time in the future over which individuals calculate their utilities to make a behavioral decision—affects behavioral change and how that in turn influences the dynamics of an epidemic.

We introduce the model through the lens of individuals rather than at the population level, because we find that the individualistic perspective gives a more natural interpretation of the behavioral decision analysis we discuss. Under the assumption that the population is large, well mixed, and homogeneous, we can also express the model as a system of ordinary differential equations (ODEs). We work with both versions of the model, using the ODE version for calculations, and the individual-based or agent-based model (ABM) version for thinking about the psychological features of decision making that may affect the spread of infectious diseases.

One of our key findings is that individuals choose to engage in prophylactic behavior only when the planning horizon is “just right.” If the planning horizon is set too far into the future, it is in an individual’s best interest to become infected (i.e. get it over with); if the horizon is too short, individuals dismiss the future risk of infection (i.e. live for the moment). What counts as “just right” depends on the disease in question, and we explore two hypothetical contrasting diseases, one with long recovery time and acute severity, and another with short recovery time and mild severity.

## Methods

### The SPIR Model

The SPIR model couples two sub-models: one reproducing the dynamics of the infectious disease, the *Disease Dynamics Model*, and another that determines how agents make the decision to engage in prophylactic or non-prophylactic behavior, the *Behavioral Decision Model*.

### Disease Dynamics Model

The disease dynamics model reproduces the dynamics of the infectious disease in a constant population of *N* agents. Each agent can be in one of four states: Susceptible (S), Prophylactic (P), Infectious (I), or Recovered (R). The difference between agents in states S and P is that the former engage in non-prophylactic behavior and do not implement any measure to prevent infection, while the latter adopt prophylactic behavior which decreases their probability of being infected (e.g. wearing a mask, washing hands, etc.). Agents in the infectious state I are infected and infective, while those in the recovered state R are immune to and do not transmit disease. The transition between states is captured with the state-transition diagram shown in Fig. 1. For reference, all the parameters and variables in the SPIR model are listed and defined in Table 1.

**Figure 1.**
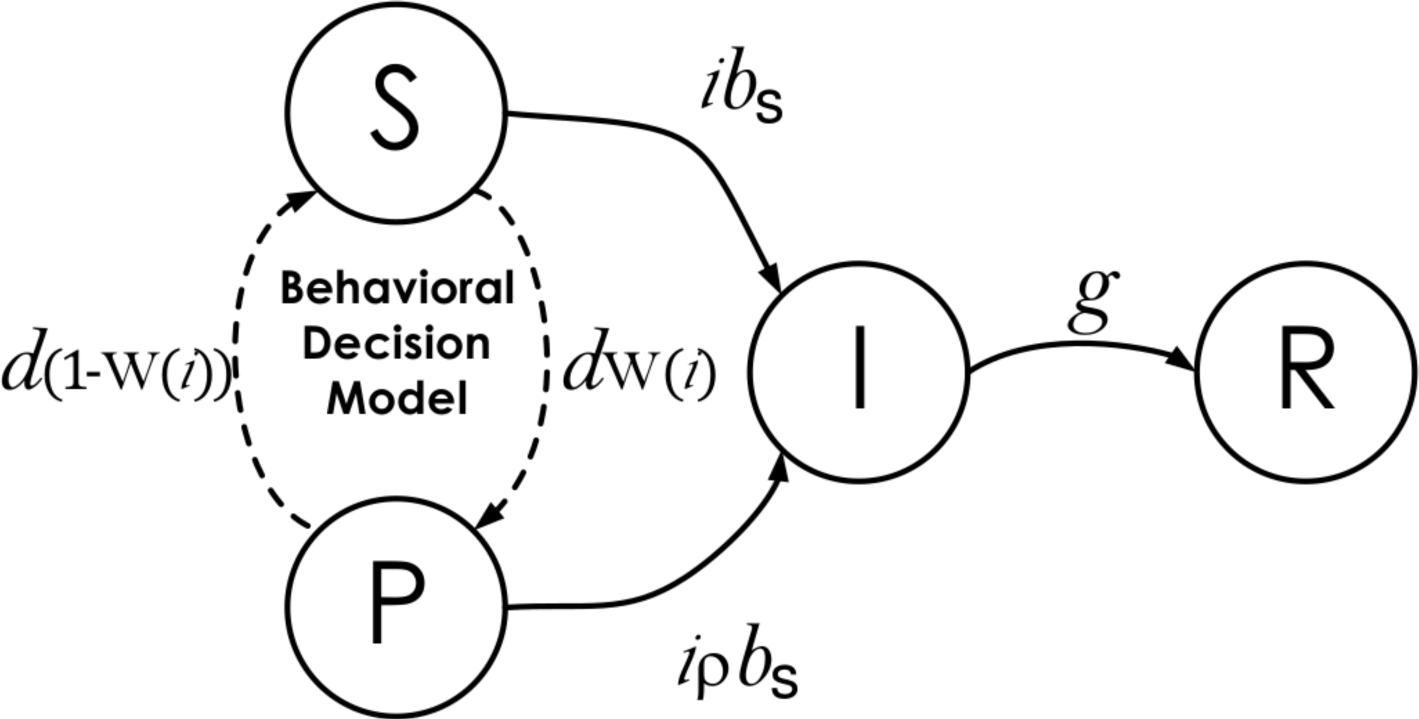
State-transition diagram of individuals in the epidemiological model. S, P, I, and R represent the four epidemiological states an agent can be in: Susceptible, Prophylactic, Infectious, and Recovered, respectively. The parameters over the transitions connecting the states represent the probability per time step that agents in one state move to an adjacent state: *i* is the proportion of infectious agents in the population; *b_S_* and *b_P_* are the respective probabilities that an agent in state S or P, encountering an infectious agent I, becomes infected; *g* is the recovery probability; *d* is the behavioral decision making probability; and W(*i*) is an indicator function returning value 1 when the utility of being prophylactic is greater than the utility of being susceptible and 0 otherwise (see details in Behavioral Decision Model).

**Table 1.**
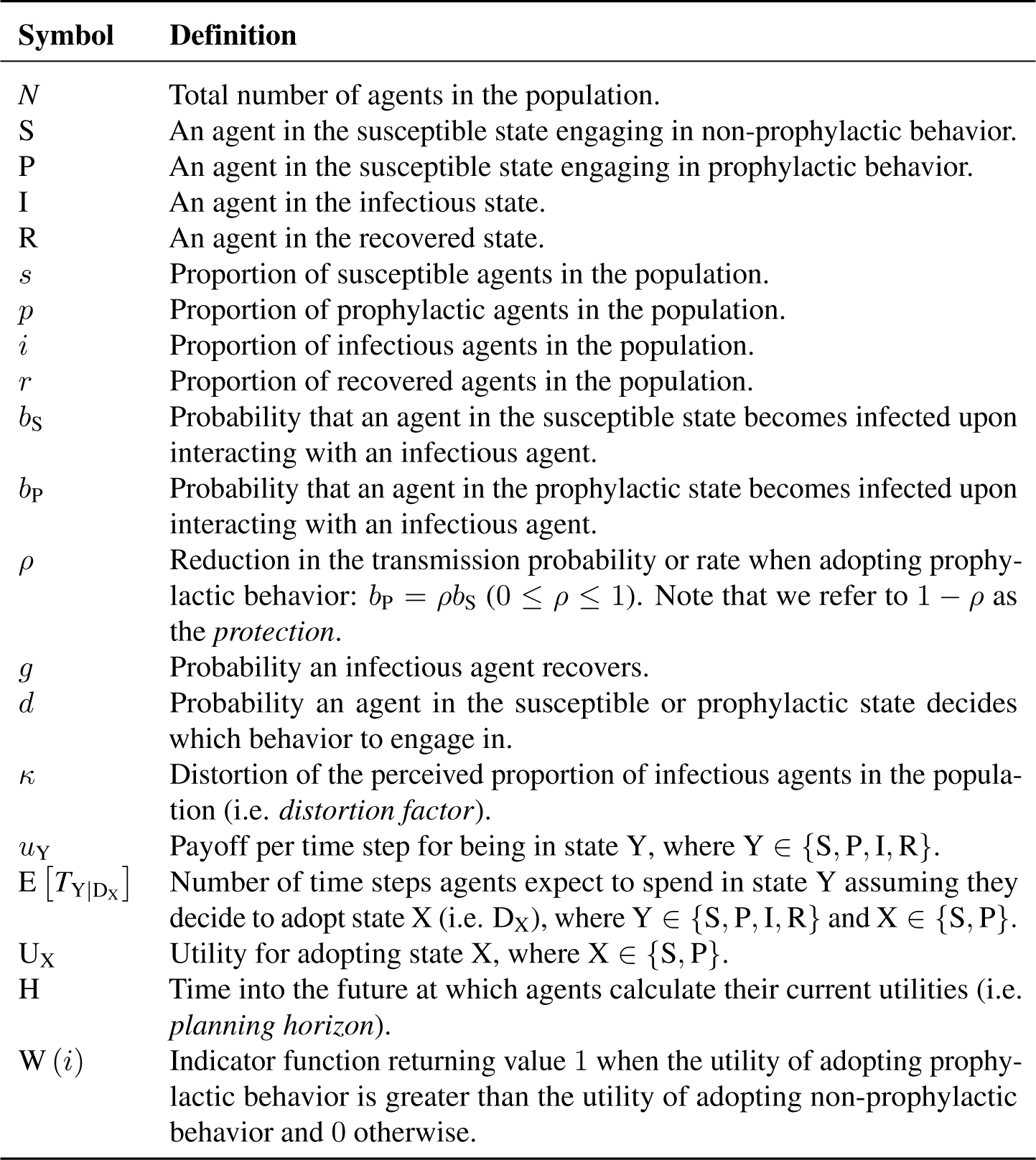
Parameters and state variables of the SPIR model.

This sub-model assumes that in each time step, three types of events occur: (i) interactions among agents and any infections that may result, (ii) behavioral decisions to engage in prophylactic or non-prophylactic behavior, and (iii) recoveries. Agents interact by pairing themselves with another randomly selected agent in the population. Given four possible states, there are ten possible pairwise interactions. However, only two types of interactions can change the state of an agent: 〈S, I〉 and 〈P, I〉. For the interaction 〈S, I〉, the susceptible agent S is infected by the infectious agent I with probability *b*_*S*_. For the interaction 〈P, I〉, the prophylactic agent P is infected by the infectious agent I with probability *b*_*P*_, where *b*_*P*_ ≤ b_S_. The probability *b*_*P*_ is linearly related to probability *b*_*S*_ by the coefficient *ρ* (i.e. *b*_*P*_=*ρb*_S_). Thus 1 – *ρ* is the protection acquired by adopting prophylactic behavior. Assuming well-mixed interactions, the proportion of infectious agents *i* represents the probability that an agent is paired with an infectious agent and the per time step probability of an agent in either state S or P being infected is either *ib*_S_ or *ib*_P_, respectively. In addition to interacting, susceptible and prophylactic agents have probability *d* per time step of making a behavioral decision to engage in prophylactic or non-prophylactic behavior. The agents’ behavioral decision is reflected in the indicator function W(*i*)(See details in Behavior Decision Model), and they engage in the prophylactic behavior when W(*i*) = 1 (i.e. adopt state P) and the non-prophylactic behavior when W(*i*) = 0 (i.e. adopt state S). Infectious agents have probability *g* per time step of recovering. We implemented an agent-based version of the disease dynamics model using the Gillespie algorithm [12] (See Supplemental Information).

If we assume that the population is well-mixed, the dynamics can be generated using a system of ODEs:

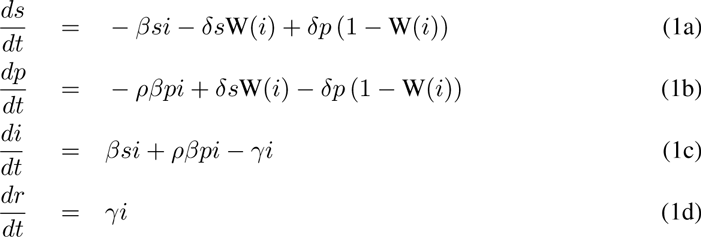

where *s, p, i*, and *r* are the proportion of susceptible, prophylactic, infectious, and recovered agents in the population. The parameters *β*, γ, and *δ* are transmission, recovery, and decision rates, whose equivalent probabilities are, respectively, transmission (*b*_S_), recovery (*g*), and decision (*d*) (Table 2). The parameter *ρ* refers to the reduction in transmission rate when adopting prophylactic behavior. We convert between rates and probabilities using equations *x* = − ln (1 − *y*) and *y* =1 − *e*^−*x*^, where *x* and *y* are rate and probability values respectively [8]. One unit of continuous time in ODE corresponds to *N* time steps in ABM.

**Table 2.**
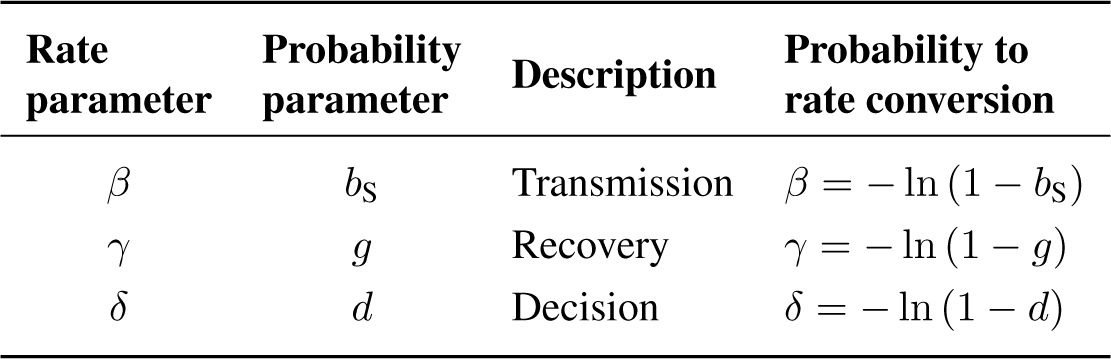
Disease parameters in rates and probabilities.

| Rate parameter | Probability parameter | Description | Probability to rate conversion |
| --- | --- | --- | --- |
| $\beta$ | $b_S$ | Transmission | $\beta = -\ln(1 - b_S)$ |
| $\gamma$ | $g$ | Recovery | $\gamma = -\ln(1 - g)$ |
| $\delta$ | $d$ | Decision | $\delta = -\ln(1 - d)$ |

### Behavioral Decision Model

Recall that agents have per step probability *d* of making a behavioral decision; here we specify how those decisions are made. The behavioral decision model, in principle, can be any model that enables agents to decide whether or not to engage in prophylactic behavior. Our decision model is a rational choice model that assumes agents are self-interested and rational; thus they adopt the behavior with the largest utility over the planning horizon, H. Note that the planning horizon is a construct used to calculate utilities and it does not affect the time until an agent has an opportunity to make another decision within the disease dynamics model.

The planning horizon is the time in the future over which agents calculate their utilities. In order for the agent to make these calculations, we make the following assumptions about agents tasked with making a decision. (i) Agents have identical and complete knowledge of the relevant disease parameters, *b*_S_, *b*_P_, and *g*. (ii) Agents’ prophylactic behavior has the same protection efficacy, *ρ*. (iii) Agents believe the current prevalence of the disease is *i* (in the case of no distortion—see below). (iv) Agents assume that *i* will remain at its current value during the next H time steps. (v) Agents compute expected waiting times based on a censored geometric distribution. Specifically, they believe that they will spend the amount of time *T*_X_ in state X €{S, P}, where the time *T*_X_ has a geometric distribution with parameter *ib*_X_, censored at the planning horizon H. When their time in X is over, agents know they will move to state I where the amount of time they expect to spend in state I has a geometric distribution with parameter *g* censored at the time remaining, H − *T*_X_. When their time in state I is over, they know they will move to state R where they will remain until time H. (vi) Agents know the per time step payoff for each state *u*_Y_, where Y €{S, P, I, R}, and all agents are assumed to have the same set of payoff values. (vii) Agents calculate the utility from now until time H under the two possible behavioral decisions (DS or DP).

*Perfect knowledge of i*. To calculate the utilities when they have perfect knowledge of *i*, agents use the length of time they expect to spend in each state. We begin by deriving the expected time in state X, where X €{S, P}, and then use this result to derive the expected time in I and, finally, R. The expected time in state I is conditioned on T_*X*_ because agents are rational and they average the time they expect to spend on state I over all possible hypothetical combinations involving X and I up to H. To simplify notation, it is helpful to first define the force of infection for state X, *f*_X_: *f*_S_= *ib*_S_ and *f*_P_= *i*ρ*b*_S_. For the agent considering hypothetical futures, the planning horizon serves to censor all waiting times greater than H, giving them value H. This leads to the following probability mass function for the time spent in state X should they decide on behavior X (denoted *T*_X|D_X__),

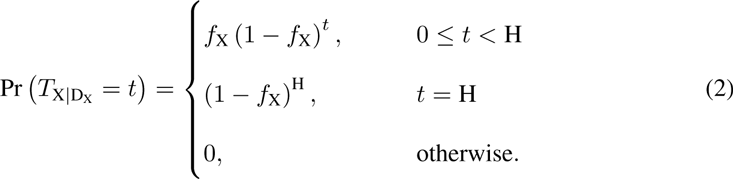

The expected time spent in state X can be expressed as

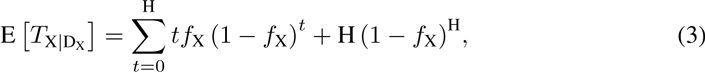

which simplifies to the desired expectation,

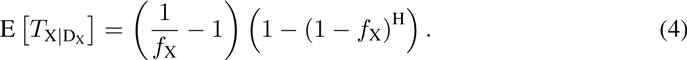

Notice that 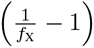 is the expected value of an uncensored geometric with minimum value of zero and that the second parenthetical term rescales the expectation to the interval [1, H].

Next, we derive the expected time spent in I by conditioning on *T*_X|D_X__.

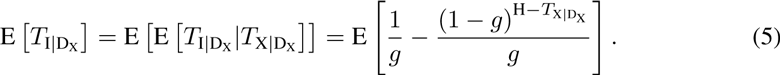

After considerable algebra, we get the expectation,

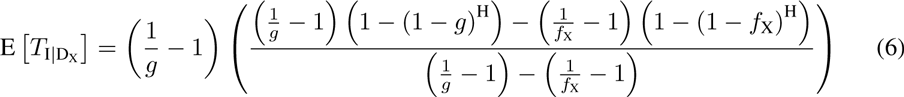

Again, the expectation of the uncensored geometric is 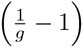. The the second parenthetical term compresses the expected time into the interval between E [*T*_X|D_X__] and H. Notice that Eq. (6) is defined only so long as *f*_χ_= *g*, When *f*_*x*_= *g*, we instead have

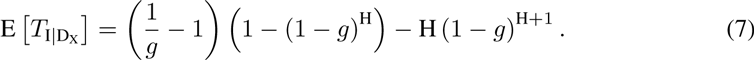

Finally, the agent calculates the expected time in state R by subtracting the expectations for X and I from H,

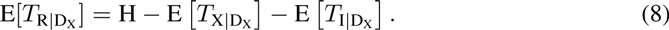

Notice that for each of the expected waiting times calculated in Eqs. (4), (6) and (7), as H goes to infinity, the rescaling terms go to one so that the equations yield the familiar expected values for the uncensored geometrics.

Having calculated these expected waiting times, the agent then calculates the utility for the two possible behaviors using,

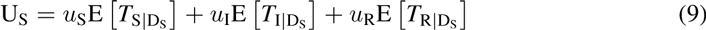

and

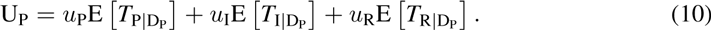

Note that when agents calculate expected times for states S and P, they need not consider the possibility of alternating to the other state in the future. This is because they assume a constant *i*which implies the best strategy now will remain the best strategy at all times during H. Thus, E[*T*_S|D_P__] and E[*T*_P|D_S__] are both zero. To be clear, this constraint pertains only to calculating utilities; agents are not constrained in how many times they actually switch states during the epidemic.

Because decisions simply reflect the largest utility and because all agents are identical, the behavioral decision can be expressed as the indicator function W (*i*) defined by

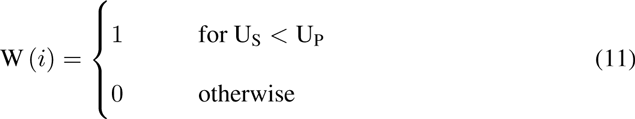

*Distorting knowledge of i*. Recall that assumption (iii) that underlies the behavioral decision model is that agents know the prevalence of the disease accurately. We relax this assumption to investigate how distorting this information effects the SPIR model. To achieve this, we replace *i* with 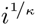 in the calculation of utilities where *κ* serves as a *distortion factor*. When *κ* = 1, *i* is not distorted; when *κ* > 1, the agent perceives *i* to be above its real value and when *κ* < 1 the opposite is true. To implement this distortion, we simply redefine *f*_X_ in the expected waiting time equations (i.e. Eqs. (3)– (7)) with 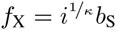 when X = S and 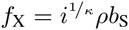 when X = P.

## Results and Analysis

The SPIR model is suitable for helping understand the influence of human behavior for diverse infectious disease epidemics. To illustrate specific characteristics of the model, however, we focus here on two contrasting diseases characterized by their severity, recovery time, and harm: *Disease 1* is acute, has a long recovery time, and may cause chronic harm, and *Disease 2* is mild, has a short recovery time, and cause no lasting harm. Table 3 shows the biological and behavioral parameter values used to generate the results discussed next, unless stated otherwise.

**Table 3.** Input parameters for two hypothetical contrasting diseases.

| Parameter |  | Disease 1 | Disease 2 |
| --- | --- | --- | --- |
| Type | Name |  |  |
| Biological | $R_0$ | 2 | 2 |
| | $\beta$ | 0.031 | 0.25 |
| | $\rho$ | 0.1 | 0.01 |
| | $\gamma$ | 0.016 | 0.125 |
| | Recovery Time ($1/\gamma$) | 65 | 8 |
| Behavioral | $\delta$ | 0 | 0 |
| | $\kappa$ | 1 | 1 |
| | $\{u_S, u_P, u_I, u_R\}$ | $\{1, 0.95, 0.1, 0.95\}$ | $\{1, 0.95, 0.6, 1\}$ |

## Behavioral Decision Analysis

Here we analyze the behavioral decision model used by the agents to decide whether or not to engage in prophylactic behavior. In particular, we are interested in identifying the level of disease prevalence above which agents would switch behavior, i.e. a *switch point*. A switch point is defined as the proportion of infectious agents beyond which it would be advantageous for an agent to switch from non-prophylactic to prophylactic behavior or vice-versa.

Figure 2D shows a heat map displaying the switch points calculated for a range of planning horizons and protection efficacies for adopting prophylactic behavior for Disease 1 〈See Fig. S1 for a heat map for Disease 2). The figure is divided into three regions—*A, B*, and *C*—that correspond to the three different utility situations illustrated in Figs. 2A, 2B, and 2C respectively.

**Figure 2.**
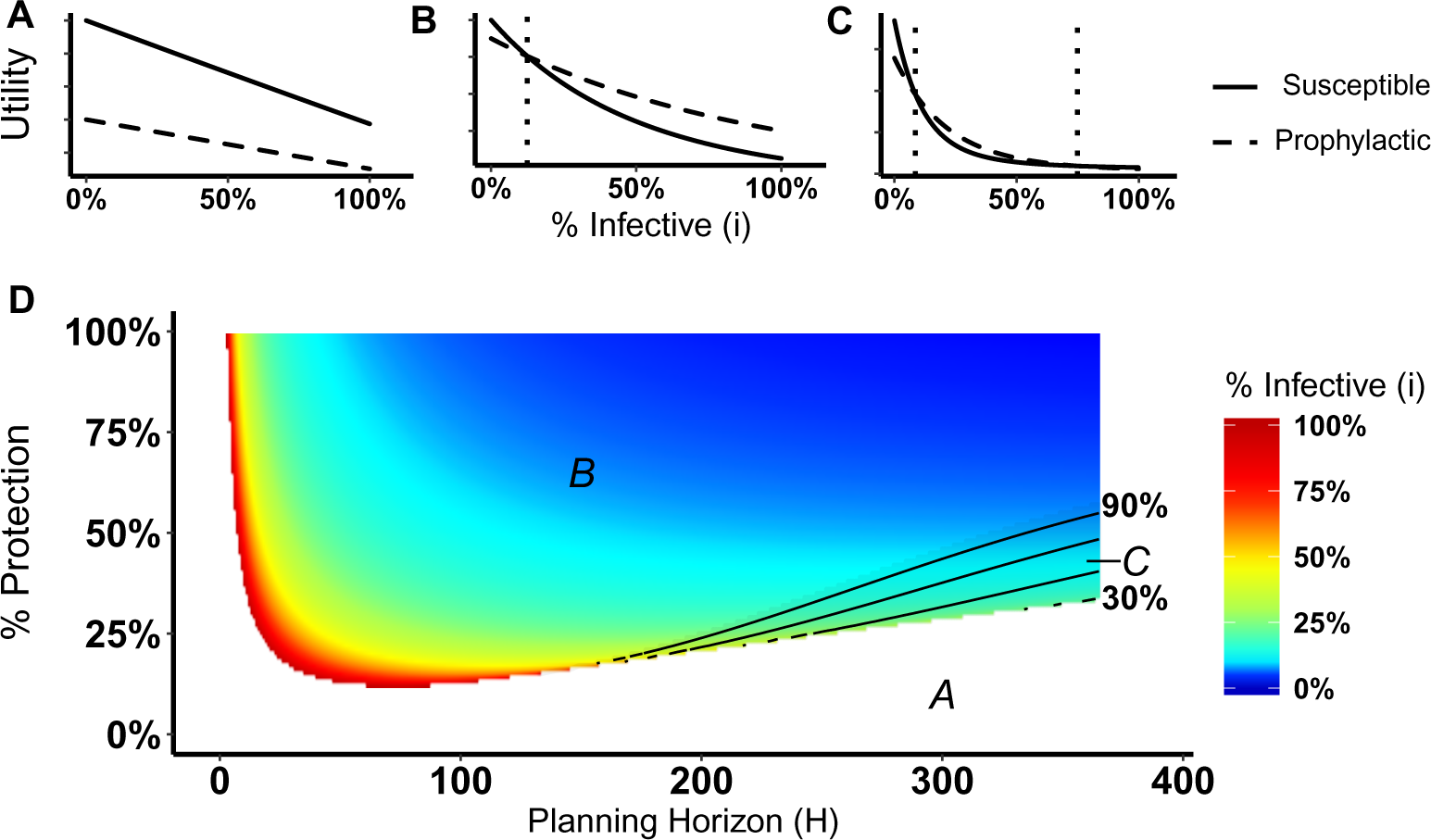
Heat map of switch points for Disease 1. (A) Situation in which non-prophylactic behavior is always more advantageous than prophylactic behavior regardless of the proportion of infectious agents. (B) Situation in which above a certain proportion of infectious agents (i.e. value indicated by the vertical dotted line), the prophylactic behavior is more advantageous than non-prophylactic behavior. (C) Situation in which prophylactic behavior is more advantageous whenever the proportion of infectious agents is within a range of values represented by the two vertical dotted lines and less advantageous otherwise. (D) Proportion of infectious agents above which prophylactic behavior is more advantageous than non-prophylactic behavior considering the percentage of protection (% Protection) obtained for adopting prophylactic behavior (1 − *ρ*) × 100 and the planning horizon H. The three regions in (D) represent the situations shown in (A), (B), and (C). In region A, agents would never adopt prophylactic behavior. In region *B*, agents would adopt prophylactic behavior above the reported proportion of infectious agents. In region *C*, agents would adopt prophylactic behavior only if the proportion of infected agents are between the proportion of infectious agents represented by the color gradient and the proportion value represented by the contour lines.

Region *A* corresponds to the situation in which agents never engage in prophylactic behavior because the utility of being in the susceptible state is never less than the prophylactic state regardless of disease prevalence (Fig. 2A). This situation occurs for low protection efficacy or short planning horizons. In the case of low protection efficacy, agents do not have an incentive to adopt prophylactic behavior because they expect to get infected regardless of their behavior. Thus, their best strategy is to become infected and then recover in order to collect the recovered payoff as quickly as possible (i.e. “get it over with”). In the case of short planning horizons, the relative contributions of the expected times of being in the susceptible or prophylactic state dominate the utilities calculation, as shown in Fig. 3. The figure illustrates how, when the planning horizon is short, the expected percentage of time spent in the susceptible or prophylactic states are greater than the expected percentage of time spent in the infectious or recovered state. Given that the susceptible payoff is greater than the prophylactic payoff, agents never adopt prophylactic behavior. The figure also shows how increasing the planning horizon changes the distribution of time spent in each state, which reduces the influence of the difference between the susceptible and prophylactic payoffs on behavioral decision.

**Figure 3.**
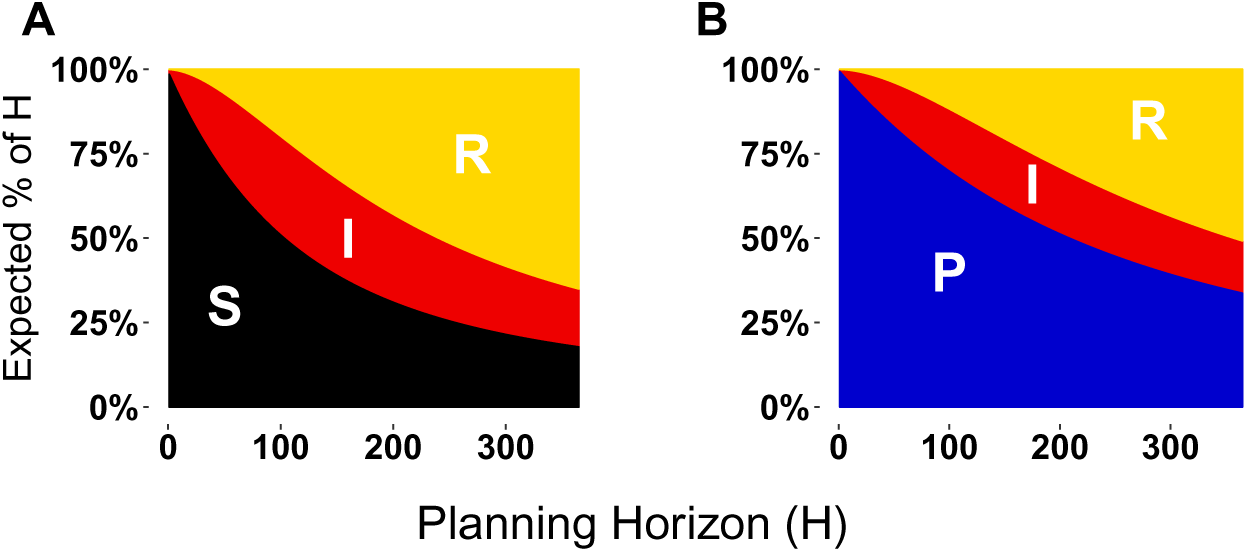
Expected proportion of the planning horizon spent in each state. (A) Proportion of the planning horizon agents expect to spend in each state if they decide to adopt state S. (B) Proportion of the planning horizon agents expect to spend in each state if they decide to adopt state P. For short planning horizons, the largest proportion of time is expected to be spent on the susceptible or prophylactic states. Increasing the planning horizon shifts the proportion of the state in which the agent will spend more time to the recovered state.

Returning our focus to Fig. 2, region *B* corresponds to the situation in which agents will adopt non-prophylactic or prophylactic behavior depending on the prevalence of the disease (Fig. 2B). If the disease prevalence is smaller than the switch point, the agent opts for the susceptible behavior and for the prophylactic behavior otherwise.

Region *C* corresponds to the situation in which two switch points exist instead of a single one (Fig. 2C). When the proportion of infectious agents is between these switch points, agents adopt prophylactic behavior, while values outside this range drives agents to adopt non-prophylactic behavior. This situation is of particular interest because it shows that the adoption of prophylactic behavior is not always monotonically associated with the prevalence of the disease.

The utility calculations that agents use to decide whether to adopt a behavior are complex (See Eqs. (9) and (10)); an exhaustive exploration of the parameter space is not undertaken here. We instead investigate several paradigm cases related to the payoff ordering. We assume that the payoff for the infectious state (*u*_I_) relies upon biological parameters of the disease and always corresponds to the lowest payoff, thus we need only consider the relationship between the other three payoffs. In particular, we are interested in looking at situations where the recovery payoff ranges from complete recovery (case 1) to less than the prophylactic state (case 4).

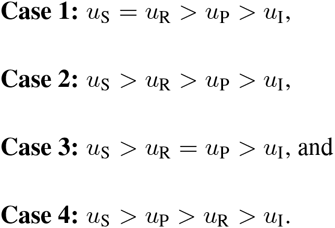

Because our model consists of a constant population of *N* agents (i.e. no mortality), cases in which *u*S > *u*R represent situations where an individual suffers chronic harm from the disease.

Figure 4 displays the switch point heat maps for these different ordering cases of Disease 1 (Figs. 4A – 4D) and Disease 2 (Figs. 4E – 4H). The most dramatic difference between the two diseases is that changing the payoff for being recovered has a large effect on the agents’ behavioral change in the cases of Disease 2, but little effect in the case of Disease 1. The reason for this has to do with the biological parameters of the model, in particular, the disease recovery time for Disease 1 is large (Recovery Time = 65), yet it is small for Disease 2 (Recovery Time =8). Consequently, an agent expects to spend more time in the recovered state when considering Disease 2 than Disease 1. When weighting these expected times with different payoffs for calculating the utilities, there will be less variation in Disease 1 compared with Disease 2.

**Figure 4.**
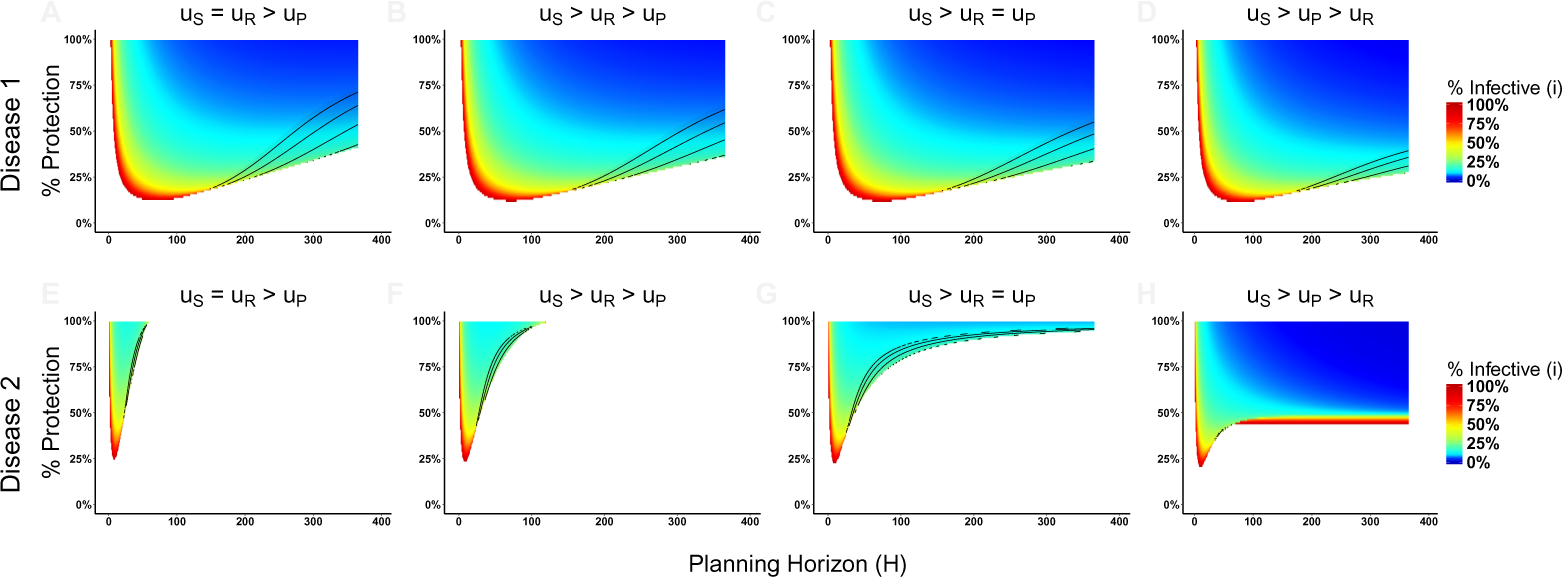
Heat maps of switch points for payoffs ordering cases of Disease 1 and Disease 2. In (A) and (E) the payoff of being susceptible and recovered are equal, which means that agents recover completely from the disease after infection. In (B) and (F), the recovered payoff is lower than the susceptible payoff, but still greater than the prophylactic payoff meaning that it is more advantageous being recovered than in the prophylactic state. In (C) and (G), any advantage comparison between being in the prophylactic or recovered state is eliminated. In (D) and (G), the disease debilitates the agent meaning that they would be better off engaging in prophylactic behavior rather be in the prophylactic state than the recovered state. The heat maps of behavioral change assume payoffs {*u*_S_, *u*_P_, *u*_I_, *u*_R_} of (A):{1, 0.95, 0.1, 1}, (B):{1, 0.95, 0.1, 0.97}, (C):{1, 0.95, 0.1, 0.95}, (D):{1, 0.95, 0.1, 0.9}, (E):{1, 0.95, 0.6, 1}, (F):{1, 0.95, 0.6, 0.97}, (G):{1, 0.95, 0.1, 0.95}, and (H):{1, 0.95, 0.1, 0.9}. The % Protection corresponds to the percentage of protection obtained for adopting prophylactic behavior (1 − *ρ*) ×100.

The effects of progressively reducing the recovered payoff are more evident for Disease 2. Reducing the recovered payoff means that lower levels of prevalence will be sufficient for agents to change their behavior. In the case of equal value for recovered and susceptible payoffs, agents consider changing behavior only in narrow parameter range of protection efficacy and planning horizon values (Fig. 4E). Progressively reducing the recovered payoff, i.e. moving from case 1 (Fig. 4E) to case 4 (Fig. 4H), the range of parameter values that would make agents change their behavior expands (i.e. there are large areas of the parameter space in which the agents would consider changing behavior) and the disease prevalence necessary for such change to occur decreases (i.e. gradual change of the color towards blue).

In addition to this numerical analysis, we have also obtained analytical results for case 2 〈Payoff ordering *u*_S_> *u*_R_> *u*_P_> *u*_I_) to identify the general conditions necessary for the existence of one or more switch points. Mathematically, switch points occur where the utility functions for S and P are equal (Eqs. (9) and (10)). Replacing the expected time notation E[*T*_Y|D_X__] in the utility *U*_S_ and *U*_P_ by the more concise 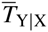, where X ∈ {S, P} and Y ∈ {S, P, I, R}, we have

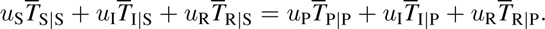

Given that 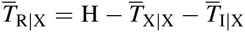,

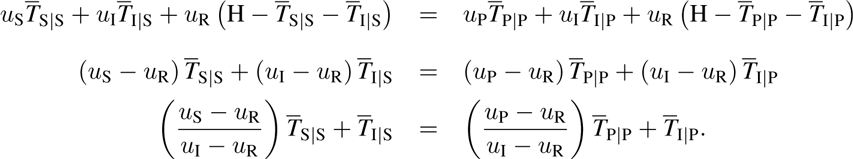

Let 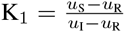, which weights the benefits of S and I, and 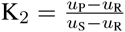, which weights the benefit of S and P. Then,

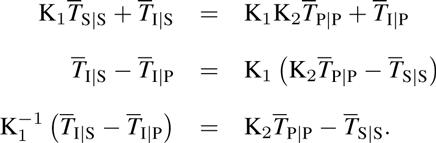

Because the payoff ordering *u*_S_> *u*_R_> *u*_P_> *u*_I_ and noting that 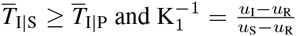, we have that 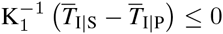 and 0 < K_2_< 1. From these we analyze some general cases.

First, we analyze the case in which 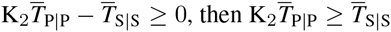. Because 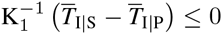, there never is a switch point and agents will strictly opt for the prophylaxis (i.e. state P).

A more interesting case is when the planning horizon H is long enough to produce the condition where 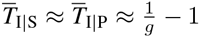 implying that

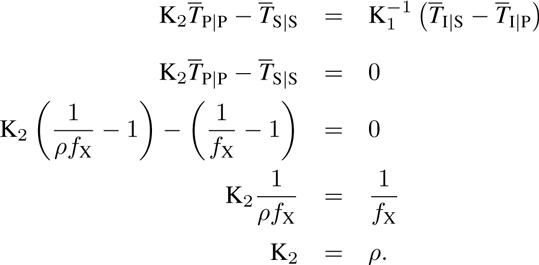

This case always produces a switch point and occurs when

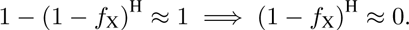

Note that if *f*_X_ is large 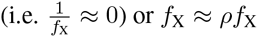), then a switch point hypothetically exists because it produces a condition where 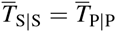. However, in these cases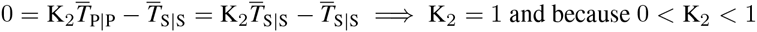, a switch point never exists.

The last case occurs when H assumes an intermediate planning horizon. When 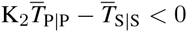, we can conclude that 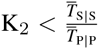 for the switch point to exist in the non-limiting case 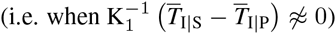. For example, if the payoff relative to recovery of P versus S is 0.5, then agents must expect less than twice the time in P to consider switching. Assuming that 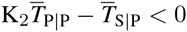, we must then further meet the condition 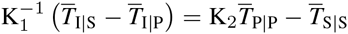 to get a switch point; this occurs when

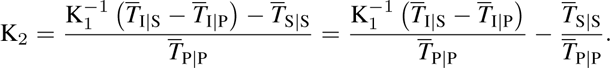

## Epidemic Dynamics

We turn now to understand how the above conditions for behavioral change may influence epidemic dynamics. Here we are particularly interested in analyzing the effects of the planning horizon H and the decision frequency d on the dynamics of Disease 1 and Disease 2.

Because we assume that the interactions among the population are well-mixed, we execute the simulations using the ODE model for a population of 100, 000 agents (initially 99, 999 agents in the susceptible state and 1 in the infectious state), decision frequency *δ* = 0.01, and protection efficacy (1 − *ρ*) = 0.9.

Figure5 shows the effects of different planning horizons on the epidemic dynamics for both Disease 1 (Fig. 5A) and Disease 2 (Fig. 5B). For short planning horizons (i.e. H = 1), agents do not ever consider changing behavior in either disease. This corresponds to the situation in Region *A* in Fig. 2A in which being prophylactic is never worth the cost, hence the epidemic dynamics are not affected. Similarly, in the cases of H = 30 for Disease 1 and H = 45 for Disease 2, we notice that neither of the two epidemic dynamics change. The dynamics are not affected because the disease prevalence does not reach the switch point (the switch points are indicated by the dashed lines in Fig. 5).

**Figure 5.**
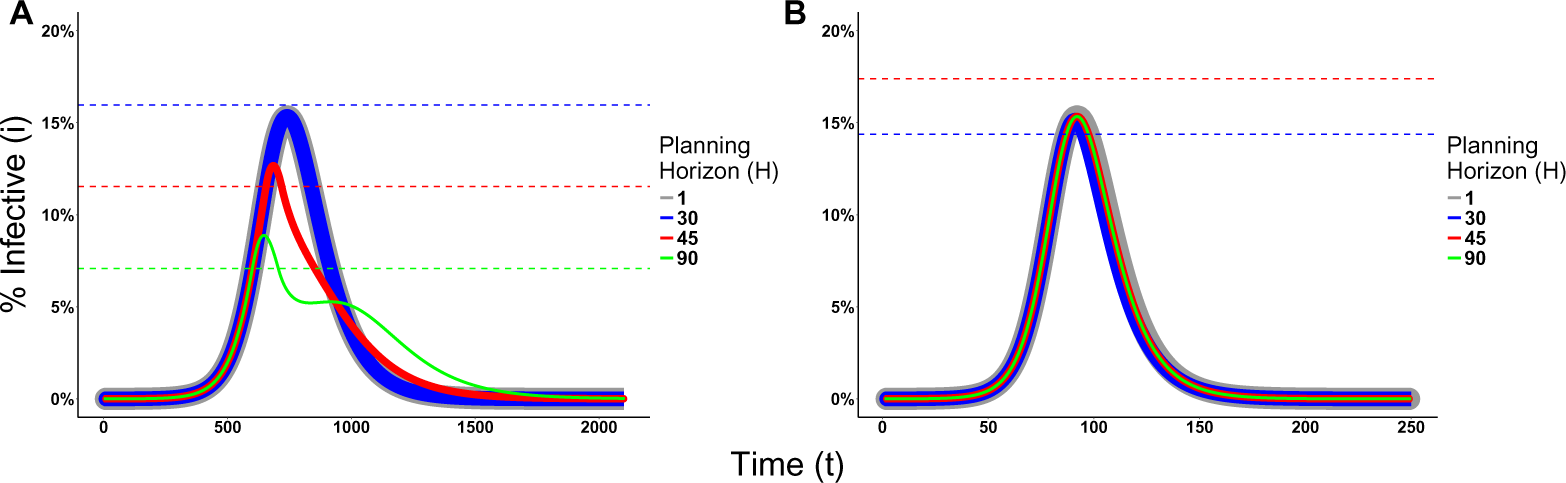
Effects of the Planning Horizon on the epidemic dynamics. (A) Disease 1 epidemic dynamics for payoffs {1, 0.95, 0.1, 0.95} and *ρ* = 0.1. (B) Disease 2 epidemic dynamics for payoffs {1, 0.95, 0.6, 1} and *ρ*= 0.01. Both dynamics consider decision frequency*δ*= 0.01. The solid lines represent the proportion of actual infectious agents in the population. The line thickness is not meaningful; rather, it is used to facilitate the visualization due to the fact that the dynamics overlap each other. The dashed lines represent the switch point associated to the planning horizon reported with the same color in the legend. Missing dashed lines indicate that no switch point exists for that planning horizon.

In the cases of H = 45 and 90 for Disease 1 and H = 30 for Disease 2, however, agents change behavior, affects the epidemic dynamics. For Disease 1, the effect is characterized by the decrease on the peak size and a prolonged duration of the epidemic. Although the dynamics of Disease 2 are also affected, the effect is small because a lower portion of the population crosses the switch point.

In other cases, increasing the planning horizon further may cause agents to never contemplate a change in their behavior, for example H = 90 for Disease 2. This means that agents willingly assume the risk of getting infected and then recover, which seems intuitive given the short recovery time and severity of the disease.

To assess the effect of the frequency that agents make behavioral decisions on the epidemic dynamics, we fix the value of the planning horizon for Disease 1 (H = 90) and Disease 2 (H = 30), and vary the decision rate. Figure 6 shows the effects of different decision frequencies on the epidemic dynamics. This figure illustrates how increasing the decision frequency reduces the peak size while prolonging the epidemic, and additionally may generate multiple waves of infection for Disease 1. These multiple waves are generated because increasing the decision frequency means individuals react faster to an increase in prevalence and adopt the prophylactic behavior. This bends the trajectory of disease incidence downward, but the reduction in prevalence causes the pendulum to swing back and individuals return back to their non-prophylactic behavior, thus creating an environment for the resurgence of the epidemic.

**Figure 6.**
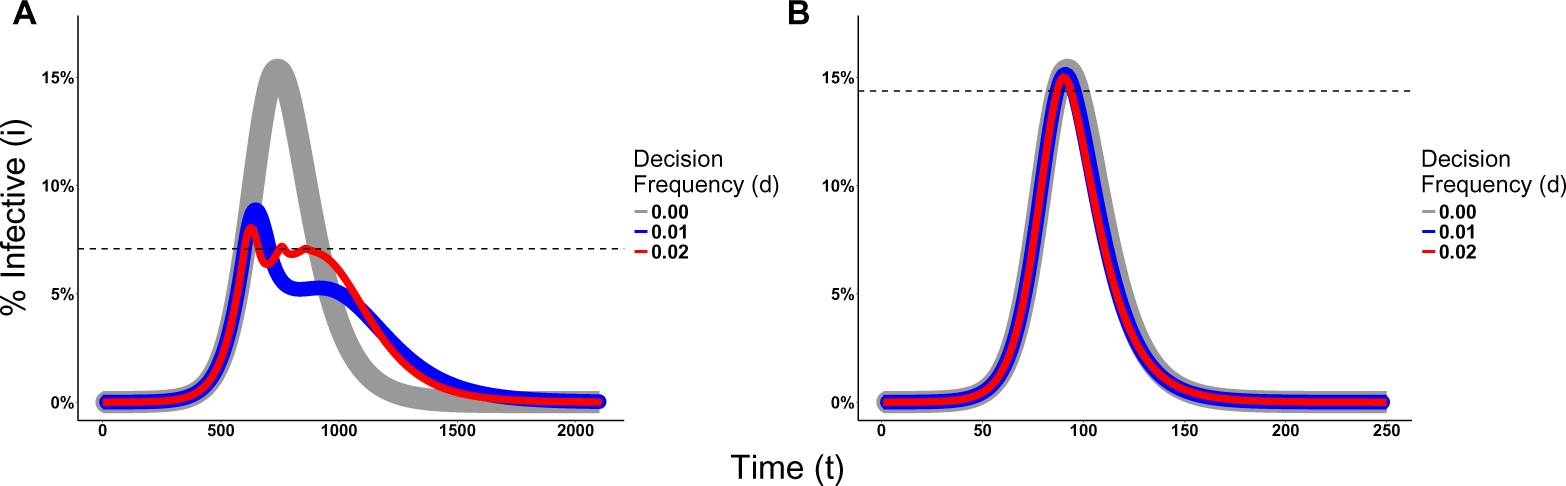
Effects of the Decision Frequency on the epidemic dynamics. (A) Disease 1 epidemic dynamics for payoffs {1, 0.95, 0.1, 0.95}, *ρ* = 0.1, and H = 90. (B) Disease 2 epidemic dynamics for payoffs {1, 0.95, 0.6, 1}, *ρ* = 0.01, and H = 30.

## Risk Perception

Empirical evidence shows that humans change behavior and adopt costly preventative measures, even if disease prevalence is low. This is especially true for harmful diseases with severe consequences to those being infected, such as Ebola or the Severe Acute Respiratory Syndrome 〈SARS). For example, despite the low level of recorded cases during the 2003 SARS outbreak in China (approximately 5, 327 cases), people in the city of Guangzhou avoided going outside or wore masks when outside [21, 20]. Combined with the severity of the disease, other factors like misinformation or excess media coverage may distort the real perception of disease prevalence (i.e. risk perception), making individuals respond unexpectedly to an epidemic.

Several models incorporate specific mechanisms regulating the diffusion of information about the disease to understand the above factors and how they contribute to the distortion of risk perception [5, 10, 18]. Here our focus is slightly different. We are interested in understanding the effects that such perception distortion has on the epidemic dynamics. Thus in our model, we have incorporated a *distortion factor κ* that alters the agents’ perception about disease prevalence used in calculating their utilities. For *κ* = 1, agents have the true perception of the disease prevalence; for *κ* > 1, the perceived disease prevalence is inflated and κreflects an increase in the risk perception of being infected; for *κ* < 1, the perceived disease prevalence is reduced below its true value.

Distorting the perception of a disease prevalence can lead to changes in the decision making process, and consequently on epidemic dynamics, as illustrated in Fig. 7 〈See Fig. S2 for Disease 2). Figure 7A shows the proportion of infectious agents above which the prophylactic behavior is more advantageous than non-prophylactic behavior assuming agents know the real disease prevalence (*κ* = 1). By distorting the perceived disease prevalence to increase the risk perception of being infected (*κ* = 1.5), the real proportion of infectious agents necessary for agents to engage in prophylactic behavior is reduced as shown in Fig. 7B. Hence, the distortion on disease prevalence makes agents engage in prophylactic behavior even when the chance of being infected is low. This affects the epidemic dynamics by reducing the peak size but prolonging the epidemic and generating multiple waves of infection as shown in Fig. 7C.

**Figure 7.**
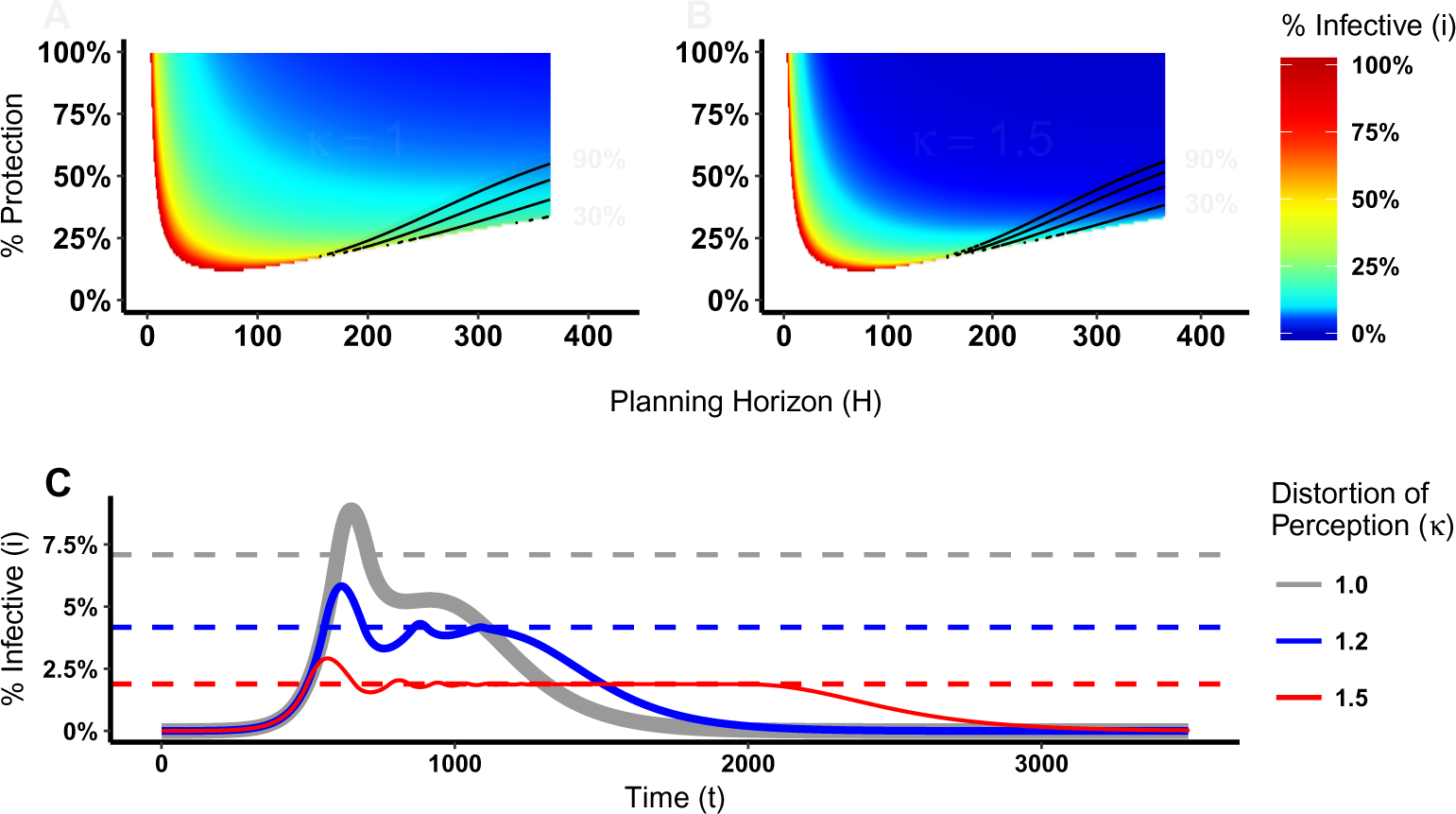
Heat maps of switch points and epidemic dynamics for Disease 1. Proportion of infectious agents above which the prophylactic behavior is more advantageous than the non-prophylactic behavior considering the percentage of protection obtained for adopting the prophylactic behavior (1 − *ρ*) × 100 and the planning horizon H. (A) No perception distortion, thus *κ* = 1; while (B) Distortion factor *κ* of 1.5, which reduces the proportion of infectious agents above which the prophylactic behavior is more advantageous. (C) Epidemic dynamics for different distortion factors that shows how increasing *κ* reduces the peak size and prolongs the epidemic.

## Discussion

Individuals acting in their own self-interest make behavioral decisions to reduce their likelihood of getting infected in response to an epidemic. We explore a decision making process that integrates the prophylaxis efficacy and the current disease prevalence with individuals’ payoffs and planning horizon to understand the conditions in which individuals adopt prophylactic behavior.

Our results show that the adoption of prophylactic behavior is sensitive to a planning horizon. Individuals with a short planning horizon (i.e. “live for the moment”) do not engage in prophylactic behavior because of its adoption costs. Individuals with a long planning horizon also fail to adopt prophylactic behavior, but for different reasons. They prefer to “get it over with” and enjoy the benefits of being recovered. In both these situations, the epidemic dynamics remain unchanged because the individuals do not have an incentive to engage in prophylactic behavior even when the disease prevalence is high. For intermediate planning horizons, however, individuals adopt prophylactic behavior depending on the disease parameters and the prophylaxis efficacy. The effects on disease dynamics include a reduction in peak size, but a prolonged epidemic.

These results are consistent with the findings of Fenichel *et al*.[6], who also concluded that behavioral change is sensitive to a planning horizon. Despite generating similar results, the SPIR and Fenichel *et al*. models differ in several aspects. In the latter, susceptible agents optimize their contact rate by balancing the expected incremental benefits and costs of additional contacts. Moreover, the agents take into consideration only the payoffs of being susceptible and recovered when optimizing the contact rates. In the SPIR model, however, agents maintain a constant contact rate, yet adopt prophylactic behavior that reduces the chance of getting infected. When agents are deciding to engage in prophylactic behavior, they take into account the payoff of all possible epidemiological states. The fact that we reach the same conclusion using different models further supports the claim that the planning horizon is a relevant decision making factor in understanding epidemic dynamics.

Although associated with the prevalence of disease, the adoption of prophylactic behavior is not always monotonically associated with it. Its adoption depends on the behavioral decision parameters. For severe diseases with long recovery time, e.g. Disease 1, the option of prophylactic behavior is less sensitive to changes in the payoffs (Fig. 4A–D) compared to less severe diseases with shorter recovery time, e.g. Disease 2 (Fig. 4E–H). Therefore, understanding the payoffs related to each disease is critical to proposing effective public policies, especially because there is not a “one-size-fits-all” solution.

Another aspect to highlight is that the beneficial adoption of prophylactic behavior can be achieved through two different public policies: change the risk perception or introduce incentives that reduce the difference between the susceptible and prophylactic payoffs. The problem with increasing the risk perception is that if it is overdone, it leads to the opposite result to the one that is desired. Because individuals perceive their risk of getting the disease as highly probable, they prefer to “get it over with” and enjoy the benefits of being recovered. In contrast, the more the prophylaxis is incentivized the better the results, e.g. reduction of epidemic peak size.

Similar to our SPIR model, Perra *et al*.[15] and Del Valle *et al*.[4] also proposed an extension to the SIR model and included a new compartment that reduces the transmission rate between the susceptible and infectious states. A clear distinction between these models and the SPIR model is that their agents do not take into account the costs associated with moving between the susceptible compartment and this new compartment. While in Perra *et al*.[15] agents make the decision to move between compartments based on the disease prevalence, in Del Valle *et al*.[4] new constant transfer rates are defined to handle the transition.

In addition to these differences, an advantage of the SPIR model with respect to all other models that implement some behavioral change is the distinction between the disease dynamics and behavioral models. This distinction renders the model flexible by making it easier to, e.g. couple other decision making processes. Consequently, this modular and flexible model architecture facilitates the execution of comparative experiments with different behavioral decision models, which we plan to perform as future work.

## Acknowledgments

Research reported in this publication was supported by the National Institute Of General Medical Sciences of the National Institutes of Health under Award Number P20GM104420. The content is solely the responsibility of the authors and does not necessarily represent the official views of the National Institutes of Health. We would like to acknowledge the support of the Institute for Bioinformatics and Evolutionary Studies Computational Resources Core sponsored by the National Institutes of Health grant number P30GM103324 that provided us computer resources to perform this study. This research made use of the resources of the High Performance Computing Center at Idaho National Laboratory, which is supported by the Office of Nuclear Energy of the U.S. Department of Energy under Contract No. DE-AC07-05ID14517.

